# Spatially resolved gene expression analysis illuminates location-specific functions in the reef-building coral *Pocillopora acuta*

**DOI:** 10.64898/2026.03.13.711677

**Authors:** Zoe Dellaert, Hollie M. Putnam

## Abstract

Reef-building coral polyps contain multiple specialized tissue types with distinct functions, from feeding and defense to symbiosis and skeleton formation. While these cell types have been characterized microscopically and more recently via single-cell RNA sequencing, spatially resolved high-throughput gene expression profiling remains limited in corals. Here we combine Laser Capture Microdissection with RNA sequencing to characterize tissue-specific gene expression in the reef building coral *Pocillopora acuta*. Oral tissues, adjacent to the seawater, exhibited 1,253 upregulated genes enriched for amino acid synthesis, transmembrane transport, signaling, environmental sensing, and secretion. These tissues showed high expression of immune and microbial-recognition genes consistent with their interface with seawater microbiota: mucins, lectins, toll-like receptors (TLRs), and *MyD88* that connects TLRs to the NF-κB pathway. Aboral tissues, which build the coral’s skeleton, exhibited 552 upregulated genes enriched for developmental processes, cell adhesion, and stimulus response. We identified strong differential expression of biomineralization- associated genes, including Chitin Synthase and Wnt pathway members, suggesting previously underdescribed roles in skeleton formation. Critically, many genes implicated in specialized functions were expressed in multiple tissues. This lack of location specificity suggests functional biomarkers will likely entail multi-gene expression patterns rather than single genes. Collectively, we highlight the need for greater spatial resolution (e.g., single cell/nuclei and spatial transcriptomics) to fully resolve coral responses within their native tissue complexity. As anthropogenic climate change increasingly threatens coral reefs, spatially resolved molecular insight into coral biology will be critical for interpreting stress response mechanisms, forecasting their limits, and applying human interventions.

## Introduction

Reef-building corals are under unprecedented threat due to anthropogenic climate change, with rising ocean temperatures driving widespread bleaching, mortality, and even the functional extinction of species in some locations [1]. Continued loss of coral species will have devastating effects, as they form the structural and ecological foundation of reef ecosystems that support marine biodiversity and sustain coastal livelihoods and food security [2–4]. The structures created by corals provide critical ecosystem services by protecting shorelines from storm-driven wave action [5] which reduces subsequent flooding-caused coastal pollution and damage [6], preventing an estimated $1.8 billion USD in annual damages in the United States alone [7]. As global temperatures continue to reach record highs [8], understanding the stress response of corals at finer biological resolution is urgently needed.

Corals are complex symbiotic meta-organisms with functionally partitioned tissue architecture, thus integrating tissue-specific processes is essential for improving predictions of responses to climate change stressors and identifying mechanisms underlying resilience. The clonal polyps that build structurally complex reef-building coral colonies may seem “simple,” but there is substantial cellular complexity within these organisms [9]. Coral cell and tissue diversity have been catalogued using observational and experimental methods (e.g., histological observation, electron microscopy, *in situ* hybridization, and immunofluorescence; reviewed in [9–14]), resulting in a wealth of knowledge over decades of work. Cnidarians have two tissue types, epidermis and gastrodermis, that surround a layer of mesoglea. This pattern occurs in two layers in mirrored symmetry, separated by the gastrovascular cavity. The body plan of each polyp can be divided into two regions generally considered above and below the gastrovascular cavity (Fig. 1): the outward-facing mouth and surrounding tentacles comprise the “oral” region, while the inward-facing and skeleton-lining tissues make up the “aboral” region. Going towards the skeleton, the tissue layers are ordered as: oral epidermis, mesoglea, oral gastrodermis, gastrovascular cavity, aboral gastrodermis, mesoglea, and aboral epidermis (i.e. calicodermis). In the oral region, the polyp interfaces directly with the external environment, sensing and responding to light, mechanical, and chemical inputs from seawater [15,16] and macro- or micro-scale organisms [17]. The oral epidermis tissue responds to these inputs differently with specialized cell types; stinging cnidocyte cells fire when triggered [16,18], mucocytes produce a protective layer of mucus [19,20], and neurons transfer information throughout the polyp, polyp-connecting coenosarc, and to neighboring polyps [15,21] (Fig. 1). In both oral and aboral gastrodermis layers, symbiocytes contain arrested lysosomes known as symbiosomes, containing corals’ key photosynthetic endosymbionts in the family Symbiodiniaceae [9,13,20,22,23]. In the aboral region, gastrodermis cells excrete digestive enzymes [20], whereas epidermal desmocytes anchor the tissue to its skeleton [24] and epidermal calicoblastic cells control the organization and growth of the calcium carbonate skeleton [25–27] (Fig. 1). While several cell types, such as calicoblastic cells, are specific to certain tissue layers, others, such as neurons and immune cells, have been characterized in both epidermal and gastrodermal layers [28,29]. Thus, what may appear as a simple mirrored tissue structure, contain complex cellular activities that require high resolution spatial examination to characterize function.

**Figure 1.**
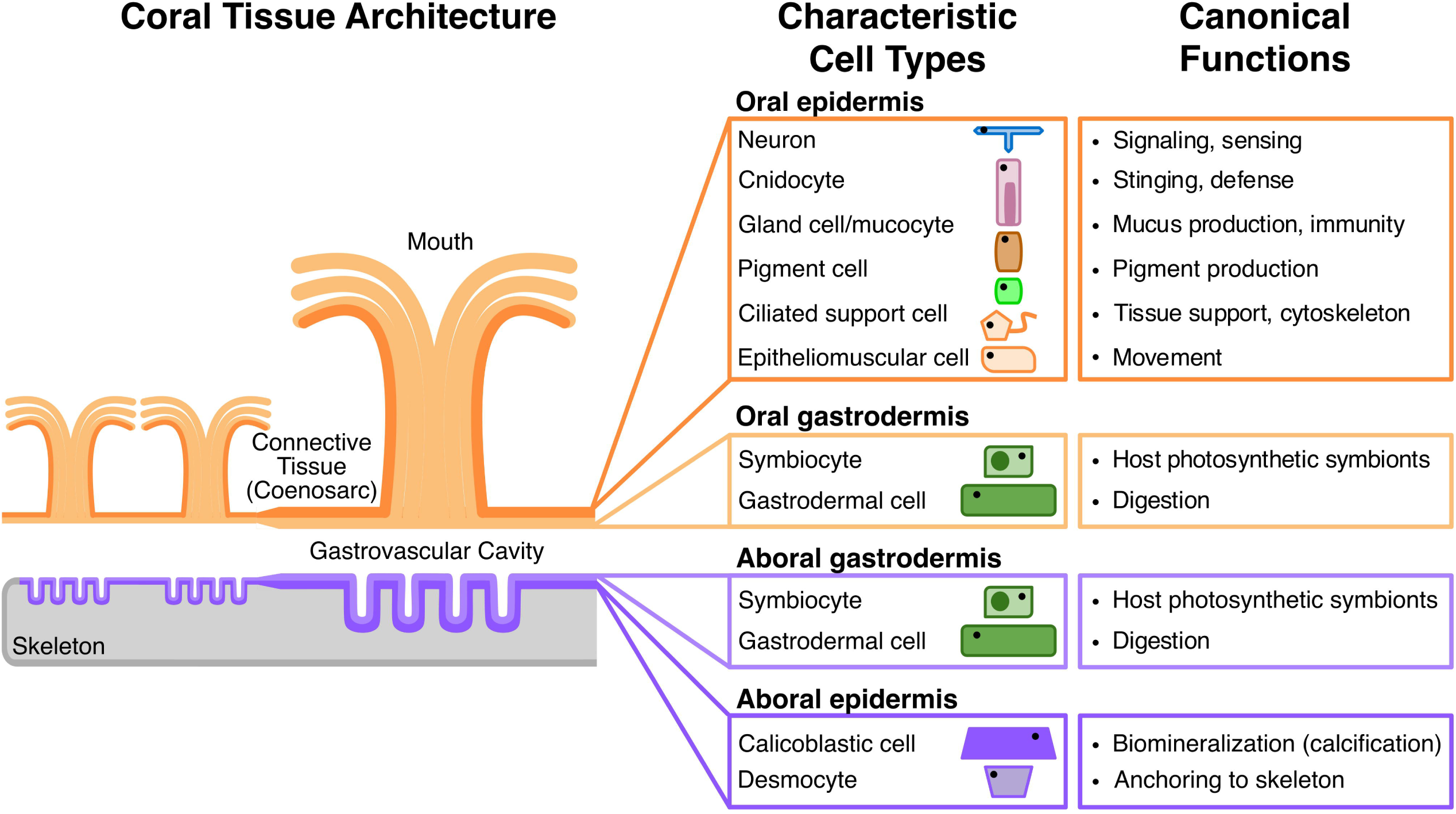
*Pocillopora acuta* coral architecture and cell types. Simplified coral tissue diagram demonstrating spatial patterning of oral and aboral epidermis and gastrodermis tissues in *P. acuta*, as well as the characteristic cell types in these locations and their canonical functions.

Histology, electron microscopy, *in situ* hybridization, and immunofluorescence studies [9–14,30] have established the foundations of coral cell and tissue biology, whereas high-throughput single cell RNA-sequencing (scRNA) now reveals the diversity of cell types present and their transcriptional signatures. By studying the entire transcriptome in each cell, scRNA-seq studies have the potential to illuminate novel or cell-specific functions and stress responses. In *Stylophora pistillata*, for example [31], scRNA-seq identified over 40 distinct cell types across three lifestages. scRNA-seq studies of non-scleractinian cnidarians [32–39] and stony corals have greatly improved our understanding of the biology of these organisms, including evolutionary adaptations to allow for facultative symbiosis [40], cellular dynamics during tissue regeneration [41], larval settlement [42], and notably given the threat of coral bleaching, gene expression profiles of symbiocytes hosting two different species of algal symbionts during heat stress [43]. However, scRNA-seq can be prohibitively costly (e.g., $2000+ per sample) [44,45], and the methods used in cell dissociation can induce stress responses that may obscure cell type identification [46–48]. Critically, sorting individual cells removes the spatial context of where these cell types are located within the coral. While the spatial locations of cell “marker” genes identified via scRNA are often verified with *in situ* hybridization, the method of scRNA inherently relies on reconstructing a picture based on highly expressed “marker” genes where spatial information is lost. Therefore, *in vivo*, high resolution and lower cost tools are poised to advance spatially explicit biological quantification.

Spatial context is a critical but often missing dimension in sequencing-based studies of reef-building corals, limiting our ability to fully understand the biological mechanisms underlying coral biology, symbiosis, and stress responses. Studies in model taxa have shown that location, not just cell type, drives expression profiles of cells and the ways they may be exposed to and respond to stress [49–52]. However, coral gene expression studies typically rely on homogenized coral tissues or mucus swabs [53,54], which obscure any gene expression diversity caused by functionally distinct microenvironments within a colony [55–57]. Even within the spatial neighborhood of one polyp, complex morphology of coral tissue enmeshed within its calcium carbonate skeleton [58] and the physiological processes [59] occurring within the coral holobiont lead to microniches of physio-chemical conditions [60], such that cell types in different areas of the tissue experience different conditions [56]. For example, the same cell type in areas of a colony or polyp with different symbiont densities will be exposed to different amounts of light [61] and oxygen [59,62]. Studies performed on tissue homogenates effectively average information across functionally distinct tissues and symbiotic compartments, leading to the loss of understanding of true biological diversity [54,56]. Studying scleractinian corals at the micro-spatial scale is particularly challenging due to their calcium carbonate skeleton, which complicates microscopical observation [63,64] and makes manual dissection of the thin tissue layers virtually impossible compared to other cnidarians [65]. This technical difficulty highlights the need for creative spatially-resolved approaches to study the biological processes in specific tissues and compartments.

Laser capture microdissection (LCM) enables isolation of specific coral tissues at high spatial resolution, advancing beyond full tissue homogenization. LCM was first developed to isolate specific cell structures of interest from cells in culture [66] and later to isolate tumors in fixed mammalian tissue [67]. In reef-building corals, LCM has been used to dissect out specific tissues for DNA sequencing of microbial 16S & metagenomics [68–70] and tissue-specific DNA methylation [71]. As a tool for gene expression studies, LCM has not been applied yet in corals. However, comparison of symbiotic vs. aposymbiotic gene expression of LCM-dissected epidermis and gastrodermis tissues in the sea anemone *Exaiptasia diaphana* [33] demonstrated that the largest magnitude of gene expression difference was between symbiotic gastrodermis and aposymbiotic gastrodermis tissues, not between epidermis and gastrodermis tissues. Therefore, LCM can provide a powerful tool for quantifying spatially-resolved biological responses in cnidarians, such as the complexity of reef building coral symbiosis, environmental sensing, and biomineralization.

In this study, we innovated LCM applications for reef building corals to examine how known biological differences between coral tissue types are reflected in their gene expression patterns. Using the common coral lab model *Pocillopora acuta*, we conducted RNA-seq following separation of oral and aboral tissues. We find that oral tissues primarily express genes involved in sensing and interacting with the external environment while aboral tissues express genes involved in tissue and skeletal development. At the same time, our spatial expression data also reveals that genes associated with canonical coral tissue-biased functions are not exclusively confined to their expected tissues. We further tested the spatial expression of *a priori* sets of genes with the spatially-specific function of biomineralization, as well as cell type marker genes indicated by scRNA-seq. The regional expression of these gene sets largely mirrored spatial expectations, but also reinforced the hypothesis that tissue-specific functions emerge from overlapping and spatially distributed gene regulation rather than strictly compartmentalized gene sets.

## Materials and Methods

### Coral species identification and husbandry

One genotype of *P. acuta* was obtained from Ocean State Aquatics in Coventry, RI in 2020, fragment, attached onto plugs, and maintained in a recirculating tank system of two 300L fiberglass coral tanks (96" L x 24" W x 12" H) and a 378L live rock tank (Rubbermaid FG424288BLA) containing artificial seawater (Fritz Reef Pro Mix Complete Marine Salt) at the University of Rhode Island from 2020-2024. The species was identified as *P. acuta* Type 5 Haplotype a (GenBank accession JX994073.1) via Sanger sequencing of the mTORF region with FATP6.1 and RORF following [72,73].

Water movement through each coral tank was created with two Magnetic Drive pumps (Danner 02512) connected by a PVC pipe to create horizontal cross-flow. Water was recirculated through a mesh filter sock (100µm Polyester Felt) within the live rock tank containing a protein skimmer (Coral Vue Technology AC20290, 250gal). Light was delivered using 4 AI Hydra 64 LED lights per tank programmed with the following 12:12 light dark cycle: 4-hour ramp up at 08:00, 4-hour hold at maximum intensity of ∼60 µmol photons m^-2^ s^-1^, PAR from 12:00-16:00, and 4-hour ramp down. Temperature (°C), salinity (psu), and pH (total scale) were measured weekdays using a digital thermometer (Fisher 15-077-8) and a pH/conductivity meter (Fisher STARA3250) equipped with conductivity (Thermo Scientific 013005MD) and pH (Mettler Toledo 51343101) probes. A Tris calibration curve was generated monthly and used to calculate pH values on the total scale from measured mV (pH) values and temperature using the package seacarb [74] in R version 4.5.2 [75]. At the time of sampling, the conditions were: 26.38 °C, 35.14 psu, and 8.1 pH (total scale). Data for 7/2022-7/2024, the two years prior to fixation, were 26.30±0.01 °C, 35.00±0.03 psu, and 8.01±0.004 pH (mean ±sem, Fig. S1).

### Fixation and Decalcification

Five fragments (labelled A-E) of the same genotype and approximate size (15 cm^2^) were selected from the aquarium (Fig. S2). A 1 cm-long sample was clipped from each fragment (time: 16:50) using wire cutters and each replicate was immediately placed into 5 mL of PAXgene Tissue Fixative (Qiagen 765312) in individual fixative containers for 24 hours at room temperature (5mL screw-cap tubes, Olympus Plastics 21-398). Then, samples were transferred to PAXgene Tissue Stabilizer (Qiagen 765512) at 4 °C for 24 hours. Samples were washed in RNase-free ice-cold Phosphate Buffered Saline (PBS, RPI P32080) to remove excess stabilizer solution and transferred to six-well plates filled with RNAse-free EDTA (0.5 M, pH 8.0, Invitrogen AM9262) for decalcification at 4 °C at 150 rpm on a rotator-shaker (Thermo Scientific 11-676-337). EDTA solution was changed daily until the opaque white skeleton was absent (3.5 days). All tools used for handling samples were sterilized with sequential washing with 10% bleach, DI water, 70% ethanol, RNAse decontaminant (Genesee 10-456), and DI water. All plastics used throughout the experiment (tubes, syringes, pipette tips) were certified RNAse free and sterile.

### Embedding and Cryosectioning

For cryoprotection, the tissues were transferred to 15 mL screw-cap tubes containing 10 mL of 15% sucrose solution in RNAse-free ice-cold PBS (Thermo Scientific Chemicals J6427022) at 4°C for 30 minutes, and then transferred to a 30% sucrose solution in RNAse-free ice-cold PBS at 4°C for 12 hours. To embed the tissues, samples were submerged in 4 °C-cooled OCT compound (Fisher 23-730-571) in a cryomold (Sakura Tissue-Tek® Cryomold®, VWR 25608-924), frozen on powered dry ice, and transferred to −80 °C.

Cryosectioning was performed over three separate days using a Leica CM3050S Cryostat at −22 °C onto Polyethylene Naphthalate (PEN) Frame Slides (4μm) (Leica 11600289). Slides were washed in RNAse decontaminant (Genesee 10-456) for 15 seconds, then washed twice in RNAse-free water, dried at 37 °C for 15 minutes, and sterilized using ultraviolet light for 30 minutes. Tools were washed as above, dried, and equilibrated to −20 °C for 30 minutes prior to cryosectioning. The inside of the cryostat, blades, and all sectioning materials were cleaned as described in [76]. A new blade (Fisher 14-070-60) was used for each sample. Four serial 10 μm-thick sections were adhered per slide, one slide per fragment. Once the final section was adhered, the sections were dehydrated for 30s with molecular-grade 100% ethanol and air dried for 15 minutes at room temperature in a 50mL screw-cap tube containing a silica gel desiccant packet and either immediately processed for microdissection or stored at −80 °C overnight and processed the next day.

### Staining and Laser Capture Microdissection

On the day of microdissection, slides stored at −80 °C were slowly equilibrated to room temperature to prevent condensation forming and damaging the tissue: slides were held sequentially for 30 minutes each at −20 °C, 4 °C, and room temperature (∼20 °C) before opening the tube. An ice-cold 1% (w/v) Cresyl Violet (Thermo Scientific Chemicals 229630050) solution in 100% ethanol was applied directly to the slides in a petri dish over dry ice for one minute using a syringe (Fisher 14-823-436) attached to a 0.45 μm filter (MilliporeSigma SLHV033RS). Stain and excess OCT were rinsed off with ice-cold 70% ethanol, followed by final dehydration in ice-cold 100% ethanol for one minute. Slides were then air dried for 1-2 minutes in a desiccant container (chamber based on [76]) immediately preceding LCM.

LCM was performed using a Leica LMD7 Microscope using the following conditions: power = 25 μJ, speed = 10, and pulse frequency = 500 Hz at 20 × magnification under brightfield illumination. Tissue was primarily dissected from coenosarc regions where all tissue layers were clearly visible (Fig. S3). Dissected tissue areas (excluding any excess PEN membrane that was also dissected) were measured from micrographs in FIJI [77]. Oral epidermis and aboral tissues were dissected into separate tubes for each of the 5 replicate fragments, resulting in a total of 10 samples (two tissue types per coral fragment) for RNA extraction and analysis. For each coral fragment, multiple microdissections (n = 6-11 per tissue type) from a single section were pooled into a single tube of oral epidermis or aboral tissue to obtain sufficient material for nucleic acid extraction. Individual dissections ranged in area from 2,076 μm^2^ to 33,084 μm^2^ (Total dissected area per fragment available in Table S1). Given a cell size ranging from 50 to 150 μm^2^, total dissection area ranged from ∼500-3000 cells per sample. Dissections were gravity-collected caps of open 0.2 mL tubes (Genesee 24-154) loaded into the tube holder supplied with the LCM microscope. Caps contained 40 μL of a freshly prepared digestion buffer (95 μL Zymo 2X Digestion Buffer (Zymo Research D3050-1-5), 95 μL RNAse-free water, and 10 μL Proteinase K (20 mg/mL), for a final Proteinase K concentration of 1 mg/mL (Zymo Research D3001-2-5)). Successful collection of tissue was verified by examining the tube caps using the microscope. The tubes were closed while still inverted, briefly centrifuged in a benchtop mini centrifuge to collect the solution, vortexed for 5-10s to lyse cells, centrifuged in the mini centrifuge again for 1 minute, and immediately placed on dry ice before storage at −80 °C.

### RNA extraction

Total RNA was extracted using the Zymo Quick-DNA/RNA Microprep Plus Kit (Zymo Research D7005). Lysed cells in digestion buffer were thawed on ice and 60 μL of additional, freshly-prepared digestion buffer (see above) was added to the 0.2 mL tube and mixed thoroughly via vortexing. The cells were digested for 15 minutes at room temperature. The entire volume was transferred to a 1.5 mL tube and centrifuged at 9,000 rcf for 3.5 minutes to pellet any cellular or PEN membrane debris. The supernatant (95 µL) was transferred to a new tube and combined with 190 µL DNA/RNA lysis buffer and transferred to a Zymo-Spin™ IC-XM Column (Zymo Research D7005). The rest of the extraction was performed as described in the manufacturer’s protocol, including the on-column DNAse I treatment. The quantity and quality of RNA were evaluated with an Agilent 4150 Tapestation Instrument (Agilent G2991BA) using the High Sensitivity RNA ScreenTape Assay (Agilent 5067-5579). For all samples, RNA quantity was too low to calculate an RNA Integrity Score (RIN), so the DV_200_ metric [78]—the percentage of RNA fragments of size greater than 200 bp—was calculated and samples with DV_200_ greater than 50% were used for library preparation.

### RNA Library Preparation and Sequencing

RNA libraries were prepared using the NEBNext® Single Cell/Low Input RNA Library Prep Kit for Illumina (New England Biolabs E6420S) in conjunction with NEBNext® Multiplex Oligos for Illumina® (New England Biolabs E7500S) according to the manufacturer’s protocol for Low Input RNA. 8 µl of RNA was used from each sample with concentrations ranging from 0.211-0.339 ng/µl for an input of 1.69-2.71 ng. During cDNA amplification, 17 PCR cycles were performed (Denaturation: 98 °C for 45s, 17 cycles of: 10s 98 °C, 15s 62 °C, and 3 min 72°C, followed by a final extension step at 72°C for 5 minutes). During the final library amplification step, 26 µl of cDNA was used for each reaction with concentrations ranging from 0.0232-0.24 ng/µl for an input of 0.6032-6.24 ng. 11 PCR cycles (Denaturation: 98 °C for 30s, 9 or 11 cycles of: 10s 98 °C and 75s 65 °C, followed by a final extension step at 65 °C for 5 minutes) were performed for libraries with less than 3 ng of cDNA input (n=3: Frag A aboral, Frag B oral epidermis, and Frag B aboral), and 9 PCR cycles were performed for all other libraries (n=7). All cleanups during the protocol were performed as directed using KAPA Pure Beads (Roche 07983271001). Libraries were sequenced using paired-end sequencing (2 x 150bp) using the Illumina NovaSeq X Plus platform at the Oklahoma Medical Research Foundation NGS Core, achieving a minimum sequencing depth of 30M reads per sample. Raw sequence reads are deposited in the SRA (BioProject PRJNA1209584).

### RNA Bioinformatic Processing

FastQC [79] and MultiQC [80] were used to assess raw read quality, inform trimming parameters, and assess the success of the trimming step [79,80]. Adapters, library preparation oligos, and low-quality bases (Quality score < 20) were trimmed from the ends of the reads using cutadapt [81]. Any sequences shorter than 20 bp post-trimming were discarded using the --minimum-length flag [81]. Trimmed and filtered reads were aligned to the *P. acuta* genome [82] using Hisat2 [83], yielding an average genome mapping rate of 81.71%. The gene models in the genome annotation only consisted of exonic regions of the genes, and did not contain untranslated regions (UTR) [82]. Due to the 3’-biased RNA-seq library preparation used, many of the reads aligned outside of annotated 3’ ends of genes in the *P. acuta* genome and therefore are not quantified as belonging to the non-UTR containing gene models of the *P. acuta* genome [82], even though these mRNAs were expressed. Therefore, in order to improve the genome annotation with our empirical expression data, following alignment, the 10 bam files were merged using Samtools [84] and used to create an extended gene annotation file using GeneExt [85] to extend gene models to capture putatively unannotated 3’ untranslated regions. Reference-guided assembly of the read alignments to the GeneExt-updated gene annotation file was performed with StringTie [86] using StringTie in coverage estimation mode (flag -e, estimating the coverage of the genes against reference genome, -G) and then the prepDE.py script of StringTie [86] was used to generate a gene count matrix. Analysis scripts are archived at Zenodo (https://doi.org/10.5281/zenodo.19006406) and maintained at github.com/zdellaert/LaserCoral.

### RNA-Seq Differential Expression Analysis

All downstream analyses were conducted using R version 4.5.2 [75]. Genes not expressed with a coverage of 10 counts in at least half of the samples (n=5), were removed using the function pOverA (0.5, 10) in the package genefilter [87], which provides a potentially conservative to rarely expressed genes, but statistically robust dataset. This filtering resulted in a total of 14,464 genes used in the differential expression analysis out of 33,730 in the *P. acuta* genome. Gene expression of filtered counts was transformed using a variance stabilizing transformation (blind=FALSE) for visualization with principal component analysis and heatmaps (VST-normalized). Differential expression analysis was performed on filtered counts using DESeq2 [88] with the design formula accounting for biological replicate and tissue type: “∼ Fragment + Tissue.” Log2 Fold-Changes (LFC) were shrunk using *lfcShrink* [88] to apply the “Approximate Posterior Estimation for generalized linear model” (apeglm) method [89] to reduce noise and improve LFC estimates in the cases of lowly expressed genes. Genes with adjusted p-value less than 0.05 and |LFC| > 1 after the *lfcShrink* function was applied were considered to be differentially expressed in the reporting of our results (Table S2).

### Improved Gene Annotation

To increase the number of annotated genes, we updated the annotation from the originally published genome [82] using BLAST+ [90,91] version 2.14.1. We performed a BLASTp search of all proteins in the *P. acuta* genome against the manually curated UniProtKB/Swiss-Prot database (released Oct. 2, 2024) [92], downloaded from the UniProt FTP server accessed Dec. 7, 2025. We generated a local BLAST database using the *makeblastbd* function, and BLASTp sequence similarity searches were performed with an e-value cutoff of 1 x 10^-5^ for a maximum of one target sequence per *P. acuta* protein (*-max_target_seqs 1 -max_hsps 1*). This resulted in 19,491 of 33,730 genes with UniProt matches, 19,047 of which had associated Gene Ontology (GO) annotations. Of the expressed 14,464 genes, 10,931 had UniProt matches and 10,654 had GO annotations compared to 9,482 in the original genome annotation.

### Functional enrichment

Gene ontology (GO) enrichment analysis was performed separately by tissue type: genes differentially expressed with higher expression in oral tissues (padj < 0.05, LFC > 1; 1253 genes, see *Results*) and those with significantly higher expression in aboral tissues (padj < 0.05, LFC < −1; 552 genes, see *Results*). The “background” set for enrichment analysis was the 10,654 expressed and GO-annotated genes. Enrichment was performed for 376 annotated of the 552 aboral-upregulated genes against this background set and for 826 annotated of the 1,253 oral-upregulated genes against this same background set. Enrichment analysis was performed using ViSEAGO [93], a wrapper of topGO [94] using the “elim” algorithm and “fisher” test statistic with a p-value cut-off of 0.01. Enrichment was performed for Biological Process (BP), Molecular Function (MF), and Cellular Component (CC) GO terms separately. Full results are available in supplementary info (Tables S3-S5). GO terms were clustered based on the Wang [95] method for semantic similarity distance calculation [93]. Clusters were aggregated into a dendrogram using *hclust* [75] using the ward.D2 method [96].

### Gene Lists of Interest

#### Biomineralization

Coral biomineralization, or the building of coral skeleton, has been an area of intense study, resulting in a list of genes termed the “biomineralization toolkit” [phrase coined by 97], which have been annotated in *Stylophora pistillata* [98] based on extensive proteomic data [99–103]. We identified orthologous gene pairs between *S. pistillata* and *P. acuta* using Broccoli v1.3 [104]. Of the published list of 124 *S. pistillata* biomineralization-related genes in [98], we identified 109 *P. acuta* orthologs, of which 70 were expressed in our dataset. This list was expanded to include additional proteins that did not have a 1:1 ortholog to the *S. pistillata* protein listed in the original published list by searching for the following key words in the annotated SwissProt protein names: “Collagen alpha”, “hemicentin”, “skeletal organic matrix”, “carbonic anhydrase”, “coadhesin”, and “Von Willebrand Factor D.” This lengthened the list to 221 genes, 182 of which were expressed in our dataset (Table S6). All genes in the list were categorized as “Acidic Proteins”, “Actin”, “Enzymes”, “Extracellular matrix/cell adhesion,” or “Uncharacterized biomineralization proteins” for visualization purposes based on the classifications in [101]. Z-scores of expression were calculated for each differentially expressed gene in this list (Table S7) based on the VST-normalized expression values and plotted using *ComplexHeatmap* [105].

#### scRNA-based cell type marker genes

The first scRNA expression atlas for a scleractinian coral was published for *S. pistillata* [31], identifying cell-type specific marker genes across 37 metacell clusters. Using orthologous gene pair analysis implemented in Broccoli v1.3 [104], we identified 414 1:1 orthologs in *P. acuta* from the set of 491 *S. pistillata* scRNA-seq cell type marker genes [31]. 238 of the 414 orthologs were expressed in our filtered dataset (Table S8). Differentially expressed marker genes (89/238 with adjusted p-value < 0.05 and |LFC| > 1) were visualized as mean VST-normalized expression within each tissue and plotted as a circularized bar plot using *circlize* [106].

## Results

### Oral and aboral tissues exhibit distinct baseline transcriptional programs

Tissue type significantly affected gene expression patterns of *P. acuta* (Fig. 2A-B). 1,805 of 14,464 genes expressed were differentially expressed by tissue type (Fig. 2C; adjusted p-value < 0.05 and |LFC| > 1; Table S2). Oral tissues exhibited 1,253 upregulated genes and aboral tissues displayed 552 genes with significantly higher expression (Fig. 2C). Functional enrichment revealed 63 BP GO terms enriched in oral-upregulated genes and 123 BP terms in aboral-upregulated genes (Fig. 3A-B; p-value < 0.01). Several of the top oral-upregulated genes (3 of the top 5) were unannotated (Fig 2C).

**Figure 2.**
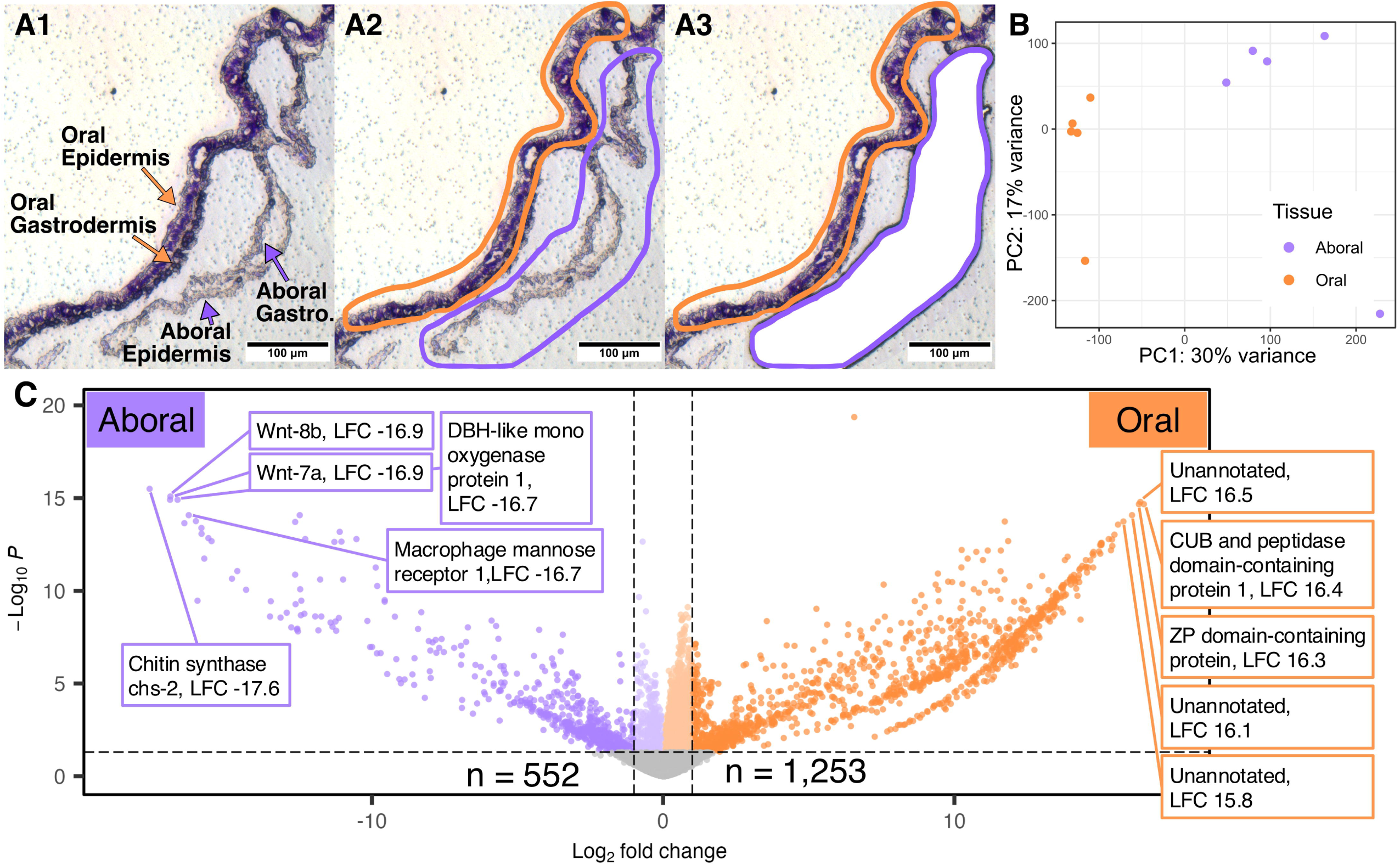
*Pocillopora acuta* displays strong tissue-specific gene expression differences. A) Laser capture microdissection of oral epidermis tissues (left, orange outline) and aboral (epidermis + gastrodermis) tissues (right, purple outline). Panel A2 shows laser path outlines; Panel A3 shows dissected aboral tissues, in the white area where the dissected slide membrane and tissue previously were. B) PCA all 14,464 expressed genes. C) Volcano plot of all expressed genes, with 1,253 differentially expressed genes (absolute value of LFC > 1 and padj < 0.05) upregulated in oral tissues shown in bold orange on the right and 552 genes upregulated in aboral tissues in bold purple on the left. Vertical dashed lines indicate the LFC cut-off used of −1 and 1, and horizontal dash lines indicate the adjusted p-value cutoff of 0.05. The top five genes by LFC are labelled.

**Figure 3.**
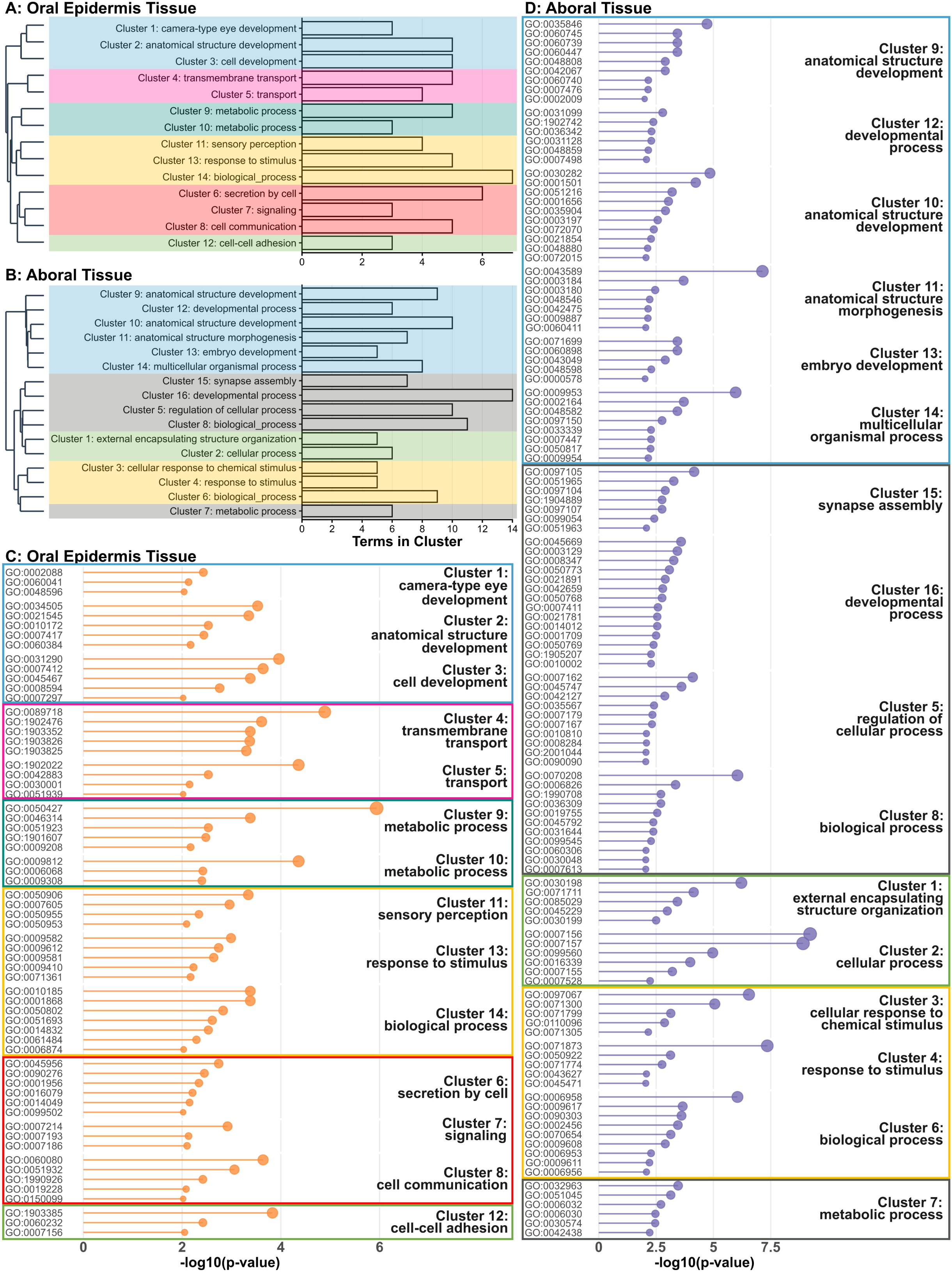
Biological Process GO Enrichment in oral and aboral tissues. A-B) GO enrichment results clustered by semantic similarity using ViSEAGO, for 14 clusters in the oral tissues (A) and 16 clusters in the aboral tissues (B). Each bar indicates the number of GO terms within a cluster enriched in genes upregulated in that tissue type. Cluster-level dendrogram of the semantic similarity clusters is included to visualize similarity between clusters. Colored boxes indicate broad functional “themes” described in the text: Blue-Development (Oral 1-3, Aboral 9-14), Pink-Transport (Oral 4-5), Teal-Metabolism (Oral 9-10), Yellow-Stimulus Response (Oral 11,13-14, Aboral 3-4, 6), Red-Signaling (Oral 6-8), Green-Cell Adhesion (Oral 12, Aboral 1-2), and Gray-Other (Aboral 5, 7-8, 15-16). C-D) Oral (C) and aboral (D) enriched GO terms ordered by p-values for each of the semantic similarity clusters.

Oral-enriched GO terms clustered into 14 semantic similarity clusters (Fig. 3C), grouping into six major functional themes: development (Clusters 1-3), transport (4-5), metabolism (9-10), signaling (6-8), stimulus response (11,13-14), and cell adhesion (12). Top enriched terms ranked by p-value were: 3’-phosphoadenosine 5’-phosphosulfate metabolic process (GO:0050427), amino acid import across plasma membrane (GO:0089718), L-lysine transport (GO:1902022), flavonoid metabolic process (GO:0009812), and retinal ganglion cell axon guidance (GO:0031290).

Aboral-enriched terms clustered into four major functional themes across 16 semantic similarity clusters (Fig. 3D): development (Clusters 9-14), cell adhesion (1-2), stimulus response (3-4, 6), and other metabolic and signaling processes (5, 7-8, 15-16). The five highest-ranked enriched BP GO terms by p-value were: homophilic cell adhesion via plasma membrane adhesion molecules (GO:0007156), heterophilic cell-cell adhesion via plasma membrane cell adhesion molecules (GO:0007157), response to norepinephrine (GO:0071873), skin morphogenesis (GO:0043589), and cellular response to thyroid hormone stimulus (GO:0097067).

### Oral epidermis tissue gene expression is enriched for amino acid metabolism and transport

Amino acid metabolic processes were significantly enriched functions in oral tissues (Fig. 3A; Oral Clusters 9-10). The top five most significantly upregulated genes contributing to these enrichments were: *PAPS synthase*, *Glutathione hydrolase 1 proenzyme*, *TabA*, *Cytokinin riboside 5’-monophosphate phosphoribohydrolase LOG3*, and *Sulfotransferase 1B1*. Two of the top five enriched BP GO terms (3’-phosphoadenosine 5’-phosphosulfate metabolic process (GO:0050427; Cluster 9) and flavonoid metabolic process (GO:0009812; Cluster 10)) were enriched because the same five Sulfotransferase genes, all upregulated in oral tissues, are the only genes in the dataset annotated with either of these GO terms. Nine genes total in the sulfation pathway were upregulated in oral tissues (*Bifunctional 3’-phosphoadenosine 5’-phosphosulfate synthase*, *Adenosine 3’-phospho 5’-phosphosulfate transporter 2*, and seven *Sulfotransferase* genes).

Transmembrane transport was also significantly enriched in oral tissues (Fig. 3A; Oral Clusters 4-5), with the most significant terms being L-lysine transport (GO:1902022; Cluster 5) and amino acid import across plasma membrane (GO:0089718; Cluster 4). The top five most significantly upregulated genes contributing to these enrichments were: *Major facilitator superfamily domain-containing protein 12*, *Monocarboxylate transporter 9 (SLC16A9)*, *Kelch-like protein 3*, *EF-hand calcium-binding domain-containing protein 4B*, and *Glycine receptor subunit alpha-1*. The biological process GO term “amino acid import across plasma membrane” (GO:0089718) was the second most significantly enriched GO term in oral epidermis tissue. All differentially expressed genes contributing to this enrichment were solute carrier (SLC) family proteins, primarily SLC1, SLC6, SLC7, and SLC16, which mediate amino acid, energy metabolite (monocarboxylates), and neurotransmitter transport.

### Oral epidermis secretion and signaling

Cell-cell signaling and synapse activity were also key functions upregulated in oral epidermis tissue (Fig. 3A; Oral Clusters 6-8). The top term in each cluster was: inhibitory postsynaptic potential (GO:0060080; Cluster 8), gamma-aminobutyric acid signaling pathway (GO:0007214; Cluster 7), and positive regulation of calcium ion-dependent exocytosis (GO:0045956; Cluster 6). The most significantly upregulated genes contributing to these enrichments were: *Synaptotagmin-10*, *Neuropeptide FF receptor 2*, *Alpha-1A adrenergic receptor*, *Adenosine receptor A2b*, and *Calcium/calmodulin-dependent protein kinase type II subunit alpha*. Of the twelve Synaptotagmin genes that were differentially expressed, ten were upregulated in oral epidermis tissue.

### Oral epidermis extracellular matrix formation and cell adhesion functions

The sixth most enriched GO term in oral epidermis tissue was regulation of homophilic cell adhesion (GO: 1903385; Cluster 12, Fig. 3A). Four differentially expressed paralogs of the cell adhesion receptor *Tyrosine-protein phosphatase Lar* were the only genes annotated with this GO term, all of which were upregulated in oral epidermis tissue. The other two enriched GO terms in Cluster 12 were delamination (GO:0060232) and homophilic cell adhesion via plasma membrane adhesion molecules (GO:0007156), with the following significantly upregulated genes contributing to their enrichments: *Hemicentin-2, Hemicentin-1, Cell adhesion molecule DSCAML1*, and two paralogs of *Protocadherin Fat 1*.

### Aboral tissues are enriched for adhesion and extracellular matrix processes consistent with biomineralization role

Cell adhesion and extracellular matrix formation were the most significantly enriched functions in the dissected aboral tissues (Fig. 3B; Aboral Clusters 1-2). The top enriched term in each cluster was: homophilic cell adhesion via plasma membrane adhesion molecules (GO:0007156; Cluster 2) and extracellular matrix organization (GO:0030198; Cluster 2). The most significantly upregulated genes contributing to these enrichments were: *Hemicentin-1*, *Receptor-type tyrosine-protein phosphatase F*, *Uromodulin*, *Collagen alpha-1(VI) chain*, and *Tyrosine kinase receptor Cad96Ca*. Fourteen *Hemicentin* genes were differentially expressed, nine of which were upregulated in aboral tissues and five of which were upregulated in oral epidermis tissue.

Given the established role of the aboral epidermis (calicoblastic cells) in skeleton formation, we examined expression of a curated set of biomineralization-associated genes from the “Biomineralization Toolkit” [98–103] and keyword searches (see Methods). Of these 182 genes, 72 were differentially expressed between tissues, with 52 upregulated in aboral tissues where biomineralization occurs (Fig. 4). We organized these biomineralization genes into functional categories: Acidic Proteins, Actin, Enzymes, Extracellular Matrix & Cell Adhesion, and Uncharacterized Biomineralization proteins. Twenty of the differentially expressed biomineralization genes were upregulated in oral tissues across these functional categories, including several paralogs of *Hemicentin*, *coadhesin*, and *collagen* as well as *Carbonic anhydrase 7*, *actin*, *CARP8*, and three uncharacterized proteins.

**Figure 4.**
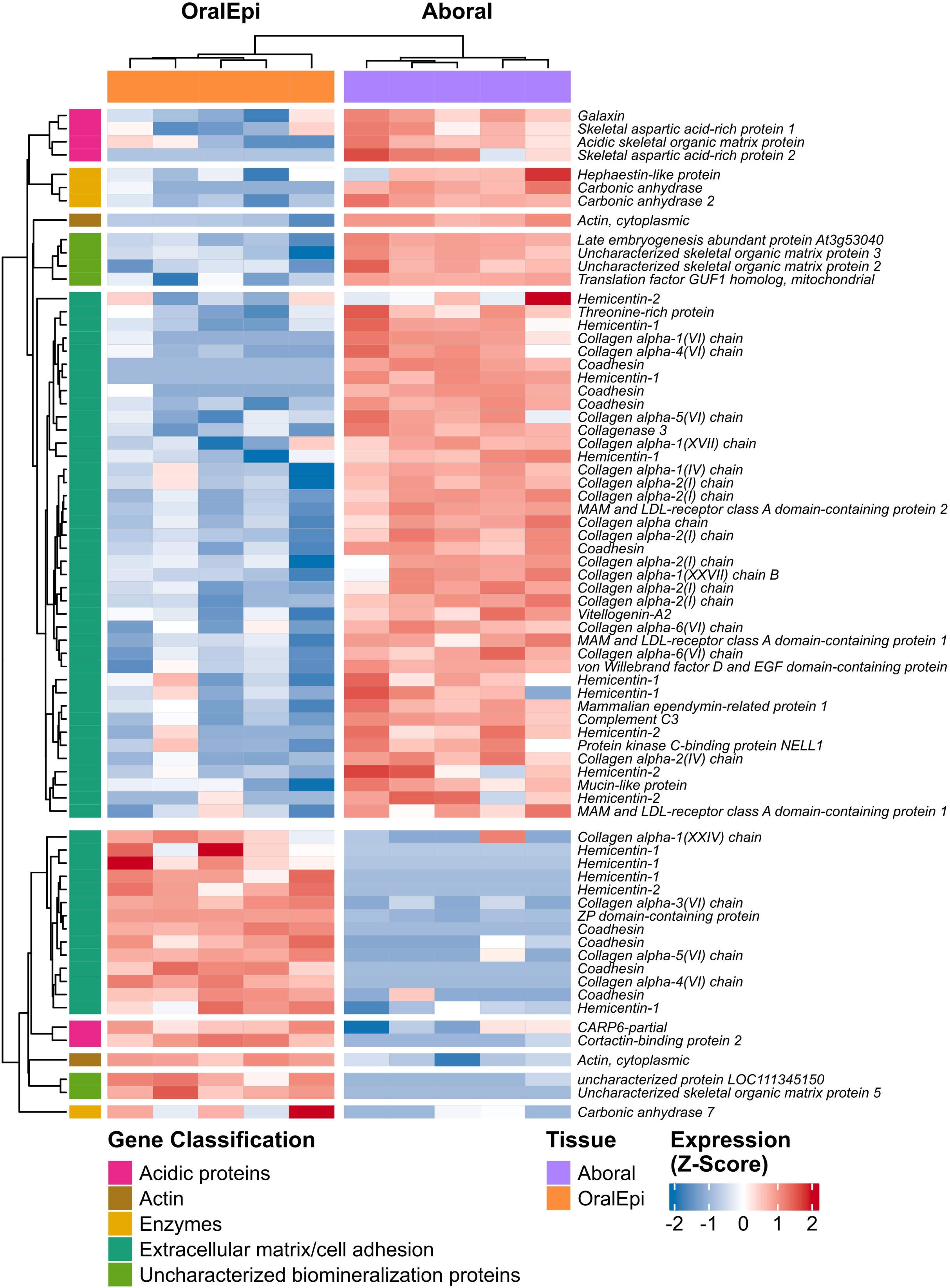
Biomineralization genes. A) Heatmap of all differentially expressed (adjusted p-value < 0.05 and |LFC| > 1) biomineralization genes from the *a priori* biomineralization gene list, categorized as “Acidic Proteins”, “Actin”, “Enzymes”, “Extracellular matrix/cell adhesion,” or “Uncharacterized biomineralization proteins”. Each column is a sample and each row of the heatmap is an individual gene, where both are grouped via k-means clustering. Rows are grouped first by the highest-order k-means cluster, which separates oral epidermis-upregulated genes and aboral-upregulated genes. Within each of those two clusters, the rows were grouped by Gene Classification, and grouped by k-means clustering within those segments. The color of each cell represents the relative expression (z-score) of that gene in that sample compared to the mean expression of that gene.

### Tissue-specific expression of developmental signaling pathways

Aboral expression was highly enriched in cell fate determination and developmental functions, spanning 66 enriched GO terms (Fig. 3B; Aboral Clusters 9-14). The top GO terms in this grouping were: skin morphogenesis (GO:0043589; Cluster 11), dorsal/ventral pattern formation (GO:0009953; Cluster 14), bone mineralization (GO:0030282; Cluster 10), oviduct epithelium development (GO:0035846; Cluster 9), and skeletal system development (GO:0001501; Cluster 10). The most significantly upregulated genes contributing to these enrichments were: *Chitin synthase chs-2*, *Wnt-8b*, *Wnt-7a*, *Receptor-type tyrosine-protein phosphatase F*, *Forkhead box protein C2-B*, and *Homeobox protein otx5-B*. Several components of the Wnt-Frizzled signaling pathway were differentially expressed, with strong upregulation in aboral tissues of *Wnt-7a*, *Wnt-8a*, and *Wnt-8b* as well as three paralogs of *Frizzled-5*a, *Secreted frizzled-related protein 5 (Sfrp1/5)*, and *Sfrp3*. Two *Wnt-2b* genes were also upregulated in oral tissues, leading to the enrichment of several developmental GO terms (Fig. 3A; Oral Clusters 1-3). The top five upregulated genes contributing to these enrichments were: *Transcription factor Sox2*, *Peroxidasin homolog pxn-2*, *Wnt-2b*, *Double-stranded RNA-specific editase 1*, and *Dynein axonemal heavy chain 5*.

### Both oral and aboral tissues respond to biotic and abiotic stimulus

Oral epidermis tissue expressed genes enriched for stimulus sensing and response (Fig. 3A; Oral Clusters 11, 13-14). The most significantly enriched terms in each cluster were: regulation of cellular defense response & regulation of complement activation, lectin pathway (GO:0010185 & GO:0001868 tied; Cluster 14), detection of stimulus involved in sensory perception (GO:0050906; Cluster 11), and detection of abiotic stimulus (GO:0009582; Cluster 13). The top five most significantly upregulated genes contributing to these enrichments were: *Beta-I spectrin*, *Retinol dehydrogenase 8*, *Neuropeptide FF receptor 2*, *Hemicentin-1*, and *Glycine receptor subunit alpha-1*.

Aboral tissues were also enriched for response to stimulus (Fig. 3B; Aboral Clusters 3-4, and 6). The most significantly enriched term in each cluster was: response to norepinephrine (GO:0071873; Cluster 4), cellular response to thyroid hormone stimulus (GO:0097067; Cluster 3), and complement activation, classical pathway (GO:0006958; Cluster 6). The most significantly upregulated genes uniquely contributing to these enrichments (i.e., not shared with Clusters 1-2 or 9-14) were: *Macrophage mannose receptor 1*, *Fibroblast growth factor receptor 3*, *Scavenger receptor cysteine-rich domain-containing protein DMBT1*, *Complement C2, Mannan-binding lectin serine protease 1*, and *Toll-like receptor 2*.

### Expression of scRNA-based cell type marker genes

To better understand how spatially-resolved dissection of tissues reflects putative cell type marker genes from scRNA-seq, and to study the distribution of cell types in our dissected tissues, we examined the expression of scRNA-seq marker genes from another Pocilloporid species: *S. pistillata* [31]. Of the 238 expressed marker genes, 89 were differentially expressed in our dataset. Forty-eight were upregulated in oral epidermis tissue and 40 were upregulated in aboral tissues (Fig. 5A). Differentially expressed *P. acuta* orthologous genes for *S. pistillata* marker genes for the following cell types were primarily upregulated in oral tissue: neuron (12 oral upregulated/14 differentially expressed), epidermis (12/12), gland (6/7), cnidocyte (3/4), immune (3/3), and germline (2/2) cell types (Fig. 5A). Gastrodermis and calicoblast cell type marker genes were primarily upregulated in aboral tissues (5/7 DE calicoblast markers and 26/28 DE gastrodermis markers; Fig. 5A).

**Figure 5.**
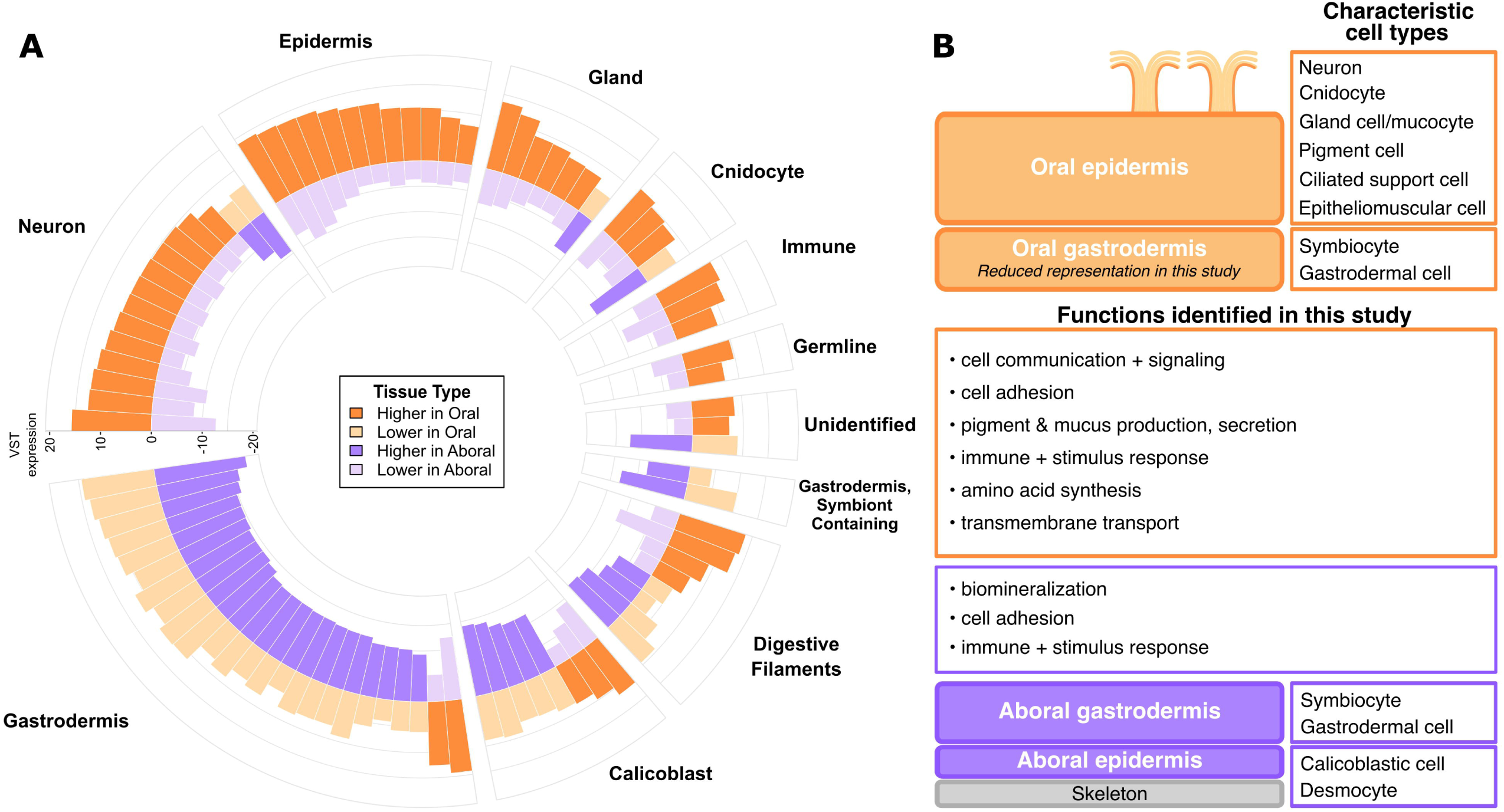
Overall patterns of tissue-specific expression of cell markers and functions. A) Circular bar plot of differentially expressed (adjusted p-value < 0.05 and |LFC| > 1) *a priori* cell type marker genes from *Stylophora pistillata* single-cell RNA-seq [31] in oral vs. aboral tissues in this study. Each sector of the circle represents a *S. pistillata* cell type, and each pair of bars (above and below the x-axis) represents the expression of an orthologous marker gene for that cell type. Each bar represents the mean VST-normalized expression level in oral samples (plotted as positive values, orange) and aboral samples (plotted as negative values, purple). Bars are colored darker to highlight the tissue of upregulation. Within each sector, bars were ordered by mean expression in the tissue with the greater number of higher expressed genes. B) Summary of functions identified in Oral and Aboral tissues, with tissue architecture and characteristic cell types depicted as in Figure 1.

## Discussion

One of the key processes in multicellular organism development is establishing tissues with distinctive locations, morphologies, and functions, a process maintained through gene expression regulation [36,107]. Though coral cell type and functional diversity have been studied for decades (reviewed in [16]), spatially resolved tissue-specific gene expression remains understudied in reef-building corals. Understanding baseline gene expression across tissues is crucial for interpreting how different tissues respond to stressors and contribute to organismal resilience, and may help characterize functions of coral “dark genes” [108,109]. In this study, we utilized LCM paired with RNA-sequencing to test whether tissue-specific gene expression patterns reflect known cellular functions in oral and aboral tissues. Importantly, we found that many genes previously implicated as markers for tissue-specific processes are actually expressed across multiple tissues, suggesting they have broader and context-dependent functions.

### Oral epidermis tissue upregulates a diverse suite of genes reflective of multiple specialized cell types

In the oral epidermis, we primarily observed functions related to amino acid synthesis and transport, cell communication, secretion, cell adhesion and extracellular matrix formation, and sensory perception and response. These observed expression patterns align with known functions of the cells located in this tissue: primarily neurons, ciliated support cells, mucus-producing mucocytes, other secretory gland cells, and pigment cells [15,16,19,20,54]. Biologically, the oral epidermis contains more diverse and specialized cell types and functions compared to the few specialized cell types in the aboral epidermis, which is potentially reflected in our data as a lack of enrichment in oral-upregulated genes. While oral tissues showed substantially more upregulated genes than aboral tissue, GO enrichment revealed relatively few oral-enriched functional categories, suggesting transcriptional diversity rather than coordination. In contrast, aboral tissues showed fewer upregulated genes, but more enriched GO terms, indicating a functionally coordinated expression profile. Notably, ten of the top twenty oral-upregulated genes lacked annotated BP GO terms, suggesting that the oral epidermis may contain cnidarian-specific or uncharacterized cell-type functions not yet captured in current functional databases.

The expression profile of the oral epidermis indicates strong sensory and neuronal activity (Figs. 3,5). The upregulation of cell communication, sensory perception-related proteins, and solute carrier (SLC) and other transport proteins, especially those involved in transport of amino acids and neurotransmitters, is consistent with elevated synaptic activity [110]. In other taxa, members of the SLC family are critical transporters of neurotransmitters at the neuron synapse [111,112], and the synaptotagmin family are key synapse-release proteins [113,114], including a calcium sensor identified to trigger synapse release [115]. The upregulation of ten *Synaptotagmin* genes, neuropeptide *LWamide* [116,117], and Sox-family transcription factors associated with neuronal and cnidocyte cell differentiation [118,119] in oral epidermis tissue in our study further support a concentration of neuronal and cnidocyte cells in this tissue region. Additionally, multiple differentially expressed genes were annotated with the GO term “stereocilium” (CC GO:0032420), all exclusively upregulated in oral epidermis tissue. The enrichment of these genes supports the presence of sensory cilia in the epidermis of cnidarians [120,121], further supporting the sensory role of the oral epidermis tissues.

The upregulation of amino acid and monocarboxylate transporters in oral epidermis tissue, including sugar transporter *SWEET1* and ammonium transporters, supports the documented role of oral tissues in nutrient exchange with symbionts [13]. However, several transporters previously implicated in facilitating coral-algal symbiosis, including SLC2 GLUT glucose transporters [31,122,123], SLC26A11 (a recently identified bicarbonate/sulfate transporter which prevents symbiosis when knocked-down in *Exaiptaisa*) [22,40], and V-type proton ATPase [123,124], were not differentially expressed between tissue types. Similarly, NPC2 (intracellular cholesterol transporter) [122], showed higher aboral expression. This pattern may reflect sampling bias: oral dissections contained primarily oral epidermis rather than gastrodermis, while aboral dissections included tightly compacted aboral gastrodermis and epidermis (Fig. 2A). Given that symbiocytes are localized primarily in the oral gastrodermis [9], this could explain the lack of strong enrichment of symbiosis-supporting genes in oral tissues (Figs. 3, 5).

Several mucus production, immune-related and microbial-recognition genes were upregulated in oral epidermis tissue consistent with this tissue’s close interactions with the external seawater and microorganisms [125–130]. Oral epidermis-upregulated genes included multiple mucin-related proteins, lectins, and toll-like receptors, as well as *MyD88*, a key protein coordinating innate immune signaling [125,131,132]. Lectins were also common in the oral epidermis-upregulated genes, consistent with their proposed role in microbial recognition by the coral host, including for recognition and uptake of its dinoflagellate symbionts [133,134]. *Nitric oxide synthase 1*, which is involved in host defense against bacteria and has also been implicated in the establishment of animal-microbe symbioses [135], was also significantly upregulated in oral epidermis tissue. While the oral epidermis tissue interfaces with the mucus layer and external microbial life, the aboral and calcifying tissue likely also require immune functions due to their lining of the gastrovascular cavity [136] as well as close contact with skeleton-associated microbes and other endolithic organisms [137]. Many immune-related genes were also upregulated in aboral tissue, as demonstrated by the enriched BP GO terms in Cluster 6, such as complement activation, classical pathway (GO:0006958; Fig. 3B). These expression patterns support the presence of coral immune cells in the oral and aboral gastrodermis tissue layers [28,29].

### Cell adhesion and extracellular matrices: key to tissue ultrastructure

Coral cells produce two key extracellular matrices: the mesoglea that lies between the epidermis and gastrodermis in both oral and aboral tissues, and the skeletal organic matrix (SOM) [138,139], both containing fibrous proteins including collagen (Fig. 4, [101,103,140–146]). Genes related to extracellular matrix production and cell adhesion were differentially expressed and enriched in both tissue types (Fig. 3). Notably, gene paralogs often showed opposing tissue-specific expression patterns, suggesting tissue-specialization of extracellular matrix components and adhesion machinery.

*Uromodulin*, a ZP-domain containing protein, exemplifies this pattern: one paralog was upregulated 15x in oral epidermis tissue, whereas another was upregulated 12x in aboral tissue. ZP-domains are recognized as “protein polymerization modules” [147] that enable proteins to polymerize into long fibrils in extracellular matrices of various kinds [148–151]. In the *P. acuta* genome, 27 genes were annotated as homologs of a ZP domain-containing protein (Uniprot G8HTB6), originally identified in the *Acropora millepora* skeletal proteome [101]. Of the 15 expressed paralogs in our dataset, 13 were significantly upregulated in oral epidermis tissue. ZP-domain containing proteins have been implicated in coral biomineralization [99–103] and spat settlement [152,153], but the expression pattern of these genes in our study supports a broader extracellular matrix (e.g., mesoglea) formation role in corals, beyond their biomineralization-associated role in the SOM.

*Hemicentin*, another extracellular matrix protein previously identified as a calicoblast marker in scRNA-seq [31], similarly showed tissue-specific differential expression. Nine of fourteen expressed paralogs were upregulated in aboral tissue, and five in oral tissue. Similar to the ZP domain-containing proteins, *Hemicentin* clearly has roles beyond biomineralization in corals, and has been documented as functioning in basement membrane-associated adhesion and extracellular matrix organization in other metazoans [154–157]. The tissue-specific expression of highly similar paralogs supports a model in which gene duplication followed by differential transcriptional regulation enables tissue-specific functions in metazoans generally [158,159] and also in cnidarian-specific biomineralization and symbiosis [65,160,161].

### Aboral gene expression reflects biomineralization and tissue organization roles

The enrichment patterns in the aboral tissues showing enrichment of adhesion and tissue development support the well-established main function of the aboral epidermis tissues: anchoring the polyp to and excreting the skeleton in the process known as biomineralization. We hypothesized that the most significantly differentially expressed genes between the two tissue types in our study would be related to this key aboral function that is wholly absent from the oral tissues. Supporting this hypothesis, the majority of a set of *a priori* biomineralization genes [‘Biomineralization Toolkit’ compiled by 98–103] were upregulated in aboral tissues (Fig. 4; Table S6), including orthologs of *Skeletal acidic Asp-rich Protein 2* and *Carbonic anhydrase*, which showed LFC in aboral tissues relative to oral tissues of ∼16x and ∼5x, respectively.

Beyond GO enrichment and this *a priori* list, though, the top upregulated genes in aboral tissues were *Chitin synthase chs-2*, *Wnt-8b*, and *Wnt-7a*. Chitin synthase, which showed the highest upregulation (∼17.5x) of all expressed genes in our dataset in aboral tissues relative to oral epidermis tissue, is the enzyme responsible for synthesizing and extracellularly secreting chitin, and is expressed in calcifying and non-calcifying cnidarians alike [162]. The role of chitin in biomineralization has not been explored much by recent studies [163–165], and as a polysaccharide it is not part of the proteomic-based “Biomineralization Toolkit” [98]. Early studies of Pocilloporid coral skeletons, however, identified a large chitinous fraction in the skeletal organic matrix [143,166]. Chitin is a known integral component of mollusc biomineralization-associated extracellular matrices, and the enzyme chitin synthase has even been hypothesized to play a key role in extracellular matrix formation by aggregating in a pH-dependent manner [167]. Chitin synthase, and chitin in general, should be further explored for their role in coral biomineralization.

The strong upregulation of Wnt pathway members in aboral tissues suggests a role in biomineralization or tissue-specific gene expression in adult corals beyond established developmental functions [168,169]. Wnt signaling controls cnidarian aboral-oral axis patterning during development, with tissue-specific Wnt expression documented in embryos [170–173]. In adult *P. acuta*, we also observed oral-aboral differences: *Wnt-7a*, *Wnt-8a*, *Wnt-8b* were upregulated in aboral tissues and *Wnt-2b* and *Wnt-2b-A* in oral tissue. This partially aligns with polyp-region patterns in the large-polyped coral *Fimbriaphyllia*, where *Wnt8* is upregulated in the body wall (epidermis and gastrodermis), but *Wnt2* and *Wnt7* are upregulated in the mouth region [174]. Colony-region expression (not tissue-specific) in adult *Acropora* [57] showed different patterns; with *Wnt2* and *Wnt8* elevated in branch tips and *Wnt7* elevated in colony bases. These patterns suggest that Wnt components maintain aboral-oral patterning and tissue organization in adults, though their specific roles remain uncharacterized in scleractinians. Critically, the divergence between adult expression patterns in our study and developmental expectations (higher oral expression in embryos) implies that non-canonical Wnt functions regulate adult tissue patterning in ways distinct from developmental programs.

Notably, the Wnt pathway interacts with the Bone Morphogenetic Protein (BMP) pathway [175,176], another key biomineralization regulator with several components also upregulated in aboral tissue. In *S. pistillata*, *BMP2/4* shows tissue-specific expression in the calicoblastic epithelium (aboral epidermis) [176], and both BMP and Wnt pathways are also upregulated during calcium supplementation-enhanced calcification [175]. In bone development, *BMP2* induces *Wnt8* expression [177], and Wnt reciprocally induces BMP expression [178,179]. This documented Wnt-BMP crosstalk in other systems [178,179], combined with their calcification-associated expression patterns in corals, may explain why Wnt upregulation in aboral tissues in our study opposes developmental expectations of higher oral expression [169,170,172,180,181].

The high expression of several hypothesized biomineralization genes in oral tissues (Table S6) reinforces that many key biomineralization proteins have roles beyond biomineralization. Carbonic anhydrases, for example, are key carbon concentrators for cnidarian-dinoflagellate symbiosis [65,182], while actin functions ubiquitously in the cytoskeleton. Ion transporters implicated in coral biomineralization, such as NKA (sodium/potassium-transporting ATPase) and NBC (SLC4-class sodium bicarbonate cotransporter), have been localized to the aboral epidermis using immunofluorescence [161,183,184] but are also found in the oral epithelia in certain species [183], suggesting multiple functions. Consistent with this, none of these genes were differentially expressed between oral and aboral tissues in our study, nor were any SLC4 bicarbonate transporters, despite their implication as critical biomineralization genes [185]. This varied or tissue-specific functionality is further supported by the finding that several proteins key to coral biomineralization are also highly expressed in non-calcifying cnidarians. For instance, SpCARP1, a coral biomineralization protein, localizes to the tentacles (oral region) of the sea anemone *N. vectensis* [186], where it functions in calcium sequestration for cellular homeostasis. This oral expression in a non-calcifying cnidarian parallels our finding of biomineralization genes in oral tissues, supporting the idea that these proteins have functions beyond skeletal formation and may even represent co-opted cellular machinery originally evolved for symbiosis, cell signaling, or other homeostatic functions [65,160,161].

### Conclusions

By combining LCM with RNA-sequencing, we revealed tissue-specific gene expression patterns in coral tissues that largely reinforce known cellular functions: the oral epidermis shows strong sensory, neuronal, and secretory activity, while the aboral tissues were dominated by biomineralization genes and tissue organization pathways. Notably, developmental signaling pathways (Wnt, BMP) show non-canonical expression patterns in adult tissues, suggesting they regulate adult tissue maintenance and biomineralization distinct from embryonic patterning. Many genes implicated in specialized functions, such as biomineralization, symbiosis, and immunity, are expressed across tissues in our study, calling into question the functional specificity of these genes. Our work confirms that different tissues have distinct baseline expression profiles, and we hypothesize they will exhibit different stress sensitivities. Future work integrating tissue- and cell-specific transcriptional responses to environmental stress, which embraces rather than averaging this complexity, is essential for predicting coral responses to climate stressors at the cellular and organismal scales.

## Acknowledgements

We would like to acknowledge Jill Ashey for library preparation troubleshooting guidance and assistance and Jill Ashey and Federica Scucchia for manuscript feedback. This work utilized resources from Unity, a collaborative, multi-institutional high-performance computing cluster managed by UMass Amherst Research Computing and Data. The sequencing at the Oklahoma Medical Research Foundation NGS Core was procured and managed via Genohub.com. The LCM instrumentation for this work was provided by Champlin grant to the College of Pharmacy and College of the Environment and Life Sciences. This work was also supported by funding from an NIH Instrumentation award 1S10OD032209-01, URI College of the Environment and Life Sciences Internal grant, and NSF Division of Biological Infrastructure grant 2316390 to HMP. This material is based upon work supported by the National Science Foundation Graduate Research Fellowship Program under Grant No 2146759. Any opinions, findings, and conclusions or recommendations expressed in this material are those of the author(s) and do not necessarily reflect the views of the National Science Foundation.

## Data Accessibility Statement

Transcriptomic Data: Raw sequence reads are deposited in the SRA (BioProject PRJNA1209584). Sample metadata: Analysis scripts and sample metadata are archived at Zenodo (https://doi.org/10.5281/zenodo.19006406) and maintained at github.com/zdellaert/LaserCoral. Sample metadata file (LCM_RNA_metadata.csv) provides Sample Name, Tissue Type, Fragment Name, Date of Cryosectioning, and Date of LCM for all samples.

## Author Contributions

ZD and HMP conceived and designed the study. ZD and HMP carried out the study and collected the data. ZD and HMP analyzed the data. ZD led the writing of the manuscript, and all authors contributed critically to interpretation and revisions. All authors gave final approval for publication.

