## Supplemental Figures and Table Captions for "Spatially resolved gene expression analysis illuminates location-specific functions in the reef-building coral *Pocillopora acuta*"

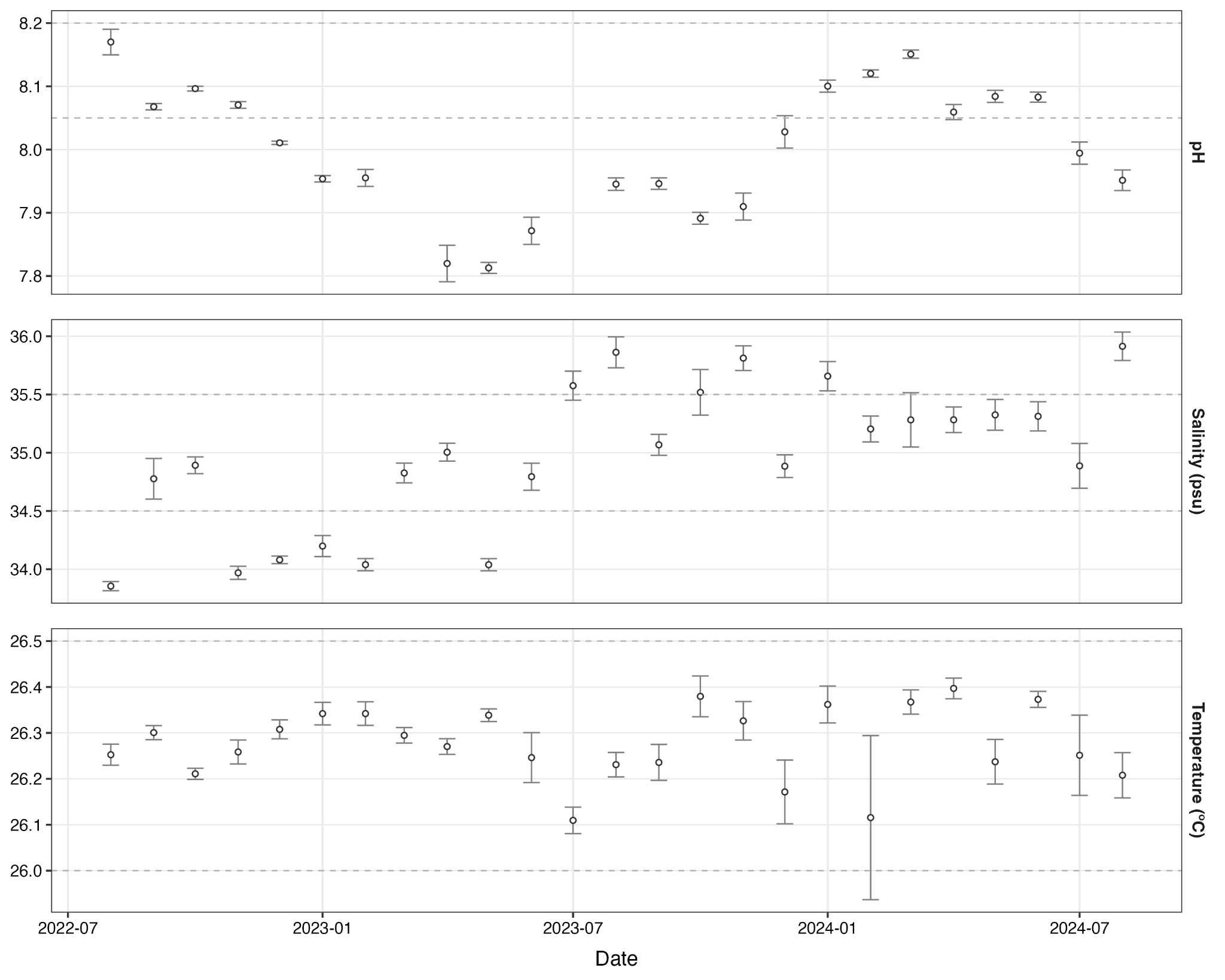

**Figure S1.** Monthly mean and standard error of Temperature (ºC), Salinity (psu), and pH (total scale) of the tank system for the two years prior to fixation.

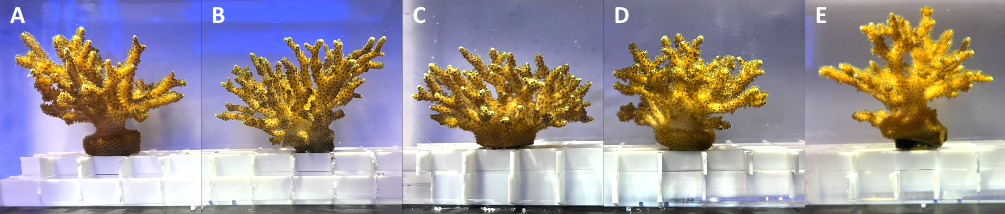

**Figure S2.** Five fragments (A-E) of *Pocillopora acuta* used in experiment.

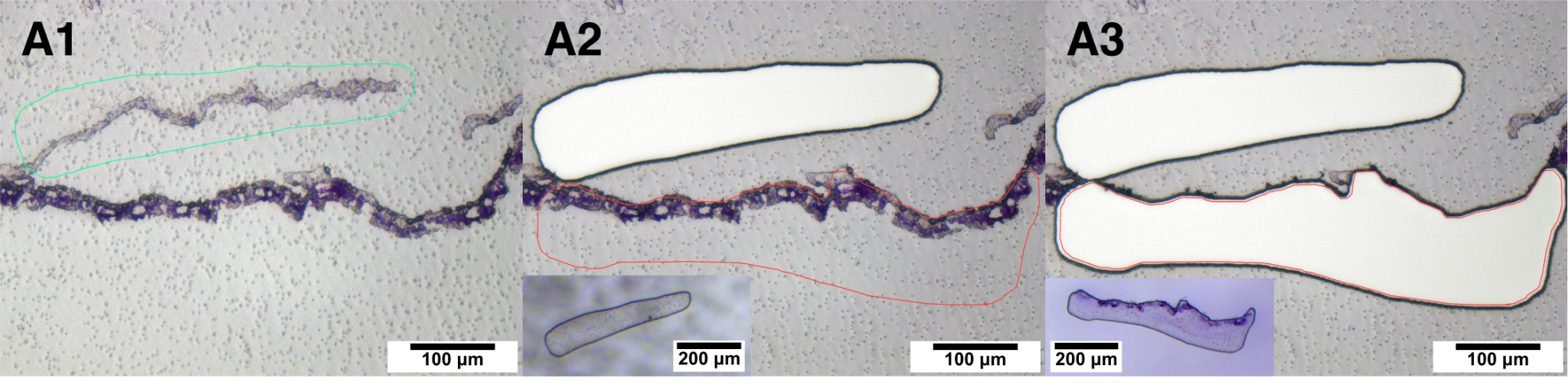

**Figure S3.** Example dissection of aboral and oral tissues in *Pocillopora acuta*. Panel A1 shows the aboral tissue dissection laser path outline (green). Panel A2 shows the oral epidermis tissue dissection laser path outline (red) and dissected aboral tissues, in the white area where the dissected slide membrane and tissue previously were. Panel A3 shows dissected oral and aboral tissues, in the white area where the dissected slide membrane and tissue previously were. Insets show the tissue + membrane collected in the tubes below the instrument.

**Table S1**. Summary of microdissected tissue areas for each coral fragment and tissue type. Multiple microdissections (n = 6-11) were pooled per sample to obtain sufficient material for nucleic acid extraction.

**Table S2.** Differential gene expression results from DESeq analysis for all expressed (14,464) genes. Reported statistics are: 1) query: *Pocillopora acuta* gene ID, 2) baseMean: the mean normalized counts across all samples, 3) log2FoldChange: Log_2_ Fold-Change between tissues, following *lfcShrink* function, 4) lfcSE: standard error of the Log_2_ Fold-Change, 5) pvalue: DESeq Wald test p-value, and 6) padj: Benjamini–Hochberg false discovery rate–adjusted p-value.

**Table S3.** Biological Process GO terms enriched in Oral and Aboral tissue, with semantic similarity-based clusters labelled. Reported data are: 1) Tissue_Upregulation: Tissue for which upregulated genes were analyzed for enrichment, 2) GO.cluster: Semantic-similarity cluster number assigned to GO term, 3) IC: information content semantic similarity score, 4) GO.ID: GO term ID number, 5) term: GO term name, 6) definition: definition of GO term, 7) elim.genes_frequency: Frequency of genes with this GO term in the test set (upregulated genes in oral or aboral tissues) compared to background (all GO-annotated genes), 8) elim.pvalue: p-value of GO enrichment test (elim algorithm, fisher test statistic), 9) elim.-log10_pvalue: log-transformed p-value, 10) elim.Significant_genes: Gene IDs of genes annotated with this GO term in the test set (upregulated genes in oral or aboral tissues), and 11) elim.Significant_genes_symbol: BLAST-annotated gene names of genes annotated with this GO term in the test set (upregulated genes in oral or aboral tissues).

**Table S4.** Molecular Function GO terms enriched in Oral and Aboral tissue, with semantic similarity-based clusters labelled. Reported data are: 1) Tissue_Upregulation: Tissue for which upregulated genes were analyzed for enrichment, 2) GO.cluster: Semantic-similarity cluster number assigned to GO term, 3) IC: information content semantic similarity score, 4) GO.ID: GO term ID number, 5) term: GO term name, 6) definition: definition of GO term, 7) elim.genes_frequency: Frequency of genes with this GO term in the test set (upregulated genes in oral or aboral tissues) compared to background (all GO-annotated genes), 8) elim.pvalue: p-value of GO enrichment test (elim algorithm, fisher test statistic), 9) elim.-log10_pvalue: log-transformed p-value, 10) elim.Significant_genes: Gene IDs of genes annotated with this GO term in the test set (upregulated genes in oral or aboral tissues), and 11) elim.Significant_genes_symbol: BLAST-annotated gene names of genes annotated with this GO term in the test set (upregulated genes in oral or aboral tissues).

**Table S5.** Cellular Component GO terms enriched in Oral and Aboral tissue, with semantic similarity-based clusters labelled. Reported data are: 1) Tissue_Upregulation: Tissue for which upregulated genes were analyzed for enrichment, 2) GO.cluster: Semantic-similarity cluster number assigned to GO term, 3) IC: information content semantic similarity score, 4) GO.ID: GO term ID number, 5) term: GO term name, 6) definition: definition of GO term, 7) elim.genes_frequency: Frequency of genes with this GO term in the test set (upregulated genes in oral or aboral tissues) compared to background (all GO-annotated genes), 8) elim.pvalue: p-value of GO enrichment test (elim algorithm, fisher test statistic), 9) elim.-log10_pvalue: log-transformed p-value, 10) elim.Significant_genes: Gene IDs of genes annotated with this GO term in the test set (upregulated genes in oral or aboral tissues), and 11) elim.Significant_genes_symbol: BLAST-annotated gene names of genes annotated with this GO term in the test set (upregulated genes in oral or aboral tissues).

**Table S6.** All expressed biomineralization genes, with annotations and references. Reported data/statistics are: 1) Heatmap_Label: Abbreviated gene name used in Figure 4, 2) query: *Pocillopora acuta* gene ID, 3) baseMean: the mean normalized counts across all samples, 4) log2FoldChange: Log_2_ Fold-Change between tissues, following *lfcShrink* function, 5) lfcSE: standard error of the Log_2_ Fold-Change, 6) pvalue: DESeq Wald test p-value, 7) padj: Benjamini–Hochberg false discovery rate–adjusted p-value, 8) List: Source of gene - either the published “biomineralization toolkit” or searched keywords (see *Methods*), 9) definition: BLAST-annotated gene name, 10) Classification: biomineralization gene classification (see *Methods*), 11) Reference: literature citation(s) for published “biomineralization toolkit” genes, and 12) Original_Accession: original accession numbers for “biomineralization toolkit” genes.

**Table S7.** Differentially expressed biomineralization genes, with z-score by sample (Figure 4). Reported data/statistics are: 1) Heatmap_Label: Abbreviated gene name used in Figure 4, 2) query: *Pocillopora acuta* gene ID, 3) baseMean: the mean normalized counts across all samples, 4) log2FoldChange: Log_2_ Fold-Change between tissues, following *lfcShrink* function, 5) lfcSE: standard error of the Log_2_ Fold-Change, 6) pvalue: DESeq Wald test p-value, 7) padj: Benjamini–Hochberg false discovery rate–adjusted p-value, 8) List: Source of gene - either the published “biomineralization toolkit” or searched keywords (see *Methods*), 9) definition: BLAST-annotated gene name, 10) Classification: biomineralization gene classification (see *Methods*), 11) Reference: literature citation(s) for published “biomineralization toolkit” genes, and 12) Original_Accession: original accession numbers for “biomineralization toolkit” genes. Remaining columns are labelled by sample names, and contain the z-score by sample for each gene (Figure 4).

**Table S8.** 238 expressed *Pocillopora acuta* orthologs of *Stylophora pistillata* scRNA-based cell type marker genes. Reported statistics are: 1) baseMean: the mean normalized counts across all samples, 2) log2FoldChange: Log_2_ Fold-Change between tissues, following *lfcShrink* function, 3) lfcSE: standard error of the Log_2_ Fold-Change, 4) pvalue: DESeq Wald test p-value, 5) padj: Benjamini–Hochberg false discovery rate–adjusted p-value, 6) Spis_CellType_Full: metacell cluster name from [[31]](https://www.zotero.org/google-docs/?nNxV8w) for which this gene was a marker, and 7) Spis_CellType: simplified metacell cluster names with numerical identifiers removed.
